# Assessment of the Genetic Diversity of Atlantic Bottlenose Dolphin (*Tursiops truncatus*) Strandings in the Mississippi Sound (USA)

**DOI:** 10.1101/2024.11.09.622766

**Authors:** Mark A. Arick, Nelmarie Landrau-Giovannetti, Chuan-Yu Hsu, Corrinne E. Grover, Stephen Reichley, Zenaida V. Magbanua, Olga Pechanova, Debra Moore, Ehsan Kayal, Anna Linhoss, Theresa Madrigal, Mark Peterman, Ozan Ozdemir, Daniel G. Peterson, Moby Solangi, Mark Lawrence, Attila Karsi

## Abstract

The common bottlenose dolphin (*Tursiops truncatus*) is a key marine mammal species in the northern Gulf of Mexico, playing an essential role as a top predator. This study focuses on the genetic diversity and population structure of bottlenose dolphins stranded in the Mississippi Sound from 2010 to 2021. A total of 511 tissue samples (muscle, liver, lung, kidney, and brain) were collected from stranded dolphins, and mitochondrial DNA (mtDNA) was extracted for analysis. Using high-throughput sequencing methods, 417 samples were successfully amplified and sequenced, producing 386 complete mitogenomes. Genetic diversity metrics, such as nucleotide and haplotype diversity, were calculated, and population structure was inferred for both mitochondrial control region (mtCR) and whole mitogenome sequences. Using the whole mitogenome, the study identified four genetically distinct populations within the Mississippi Sound, demonstrating regional variation in dolphin populations. Notably, some individuals likely originated from populations outside the sampled area. The use of whole mitogenomes allowed for improved resolution of genetic diversity and population differentiation compared to previous studies using partial mtDNA sequences. These findings provide critical insights into the genetic structure of bottlenose dolphins in the region and highlight the value of using stranded animals for population genetic studies.

## Introduction

*Tursiops truncatus* (common bottlenose dolphin) is the most abundant marine mammal species in the northern Gulf of Mexico (GoMx), occupying diverse habitats such as estuarine, coastal, continental shelf and slope, and oceanic waters[1]. They are considered essential top-level predators and a keystone species because of their impact on marine ecosystems. While common bottlenose dolphins are not considered endangered, they are a protected species in the United States under the Marine Mammal Protection Act (MMPA; 16 U.S.C 1361-1407)[2]. Under the provisions of the MMPA, the U.S. National Marine Fisheries Service (NMFS) must complete regular stock assessments of the species under its jurisdiction. They have designated 32 distinct bay, sound, and estuary (BSE) common bottlenose dolphin stocks in the northern GoMx including the Mississippi Sound, Lake Borgne, and Bay Boudreau Common Bottlenose Dolphin stock. Based on a winter 2018 aerial survey, the best-known abundance estimates for common bottlenose dolphins in this stock is 1,265 individuals[2,3].

Through photo-identification, satellite telemetry, and genetic studies, common bottlenose dolphins documented in Mississippi waters have been known to exhibit site fidelity with some groups occupying the immediate waters around the barrier islands of Mississippi (Cat Island, Ship Island, Horn Island, and Petit Bois Island) and others seasonally shifting from the inshore waters of the Mississippi Sounds to the adjacent contiguous bodies of water of the north central GoMx, Chandeleur Sound, Breton Sound, and Mobile Bay (Fig 1)[4–12]. Additionally, spatial variation and density was commonly highest in the central and eastern portions of the Mississippi Sound[6]. Because these fluctuations have been observed, further genetic studies such as the continued use of mtDNA for genetic analysis of stocks utilizing tissue collected from dead dolphins within our survey area helps in better understanding and contributing to delineating resident versus transient dolphins. Recently, Vollmer et al.[11] analyzed the mitochondrial control region (mtCR), a small but highly variable region of the mtDNA in animals, and nuclear microsatellites from bottlenose dolphin samples collected throughout the inshore waters of the Mississippi Sound and coastal waters of the north-central GoMx. The authors identified only two genetically distinct populations: one encompassing the Mississippi Sound and adjacent coastal waters (green population), the other expanding from Mobile Bay (Alabama) to the East coast of Florida (blue population)[11].

**Fig 1.**
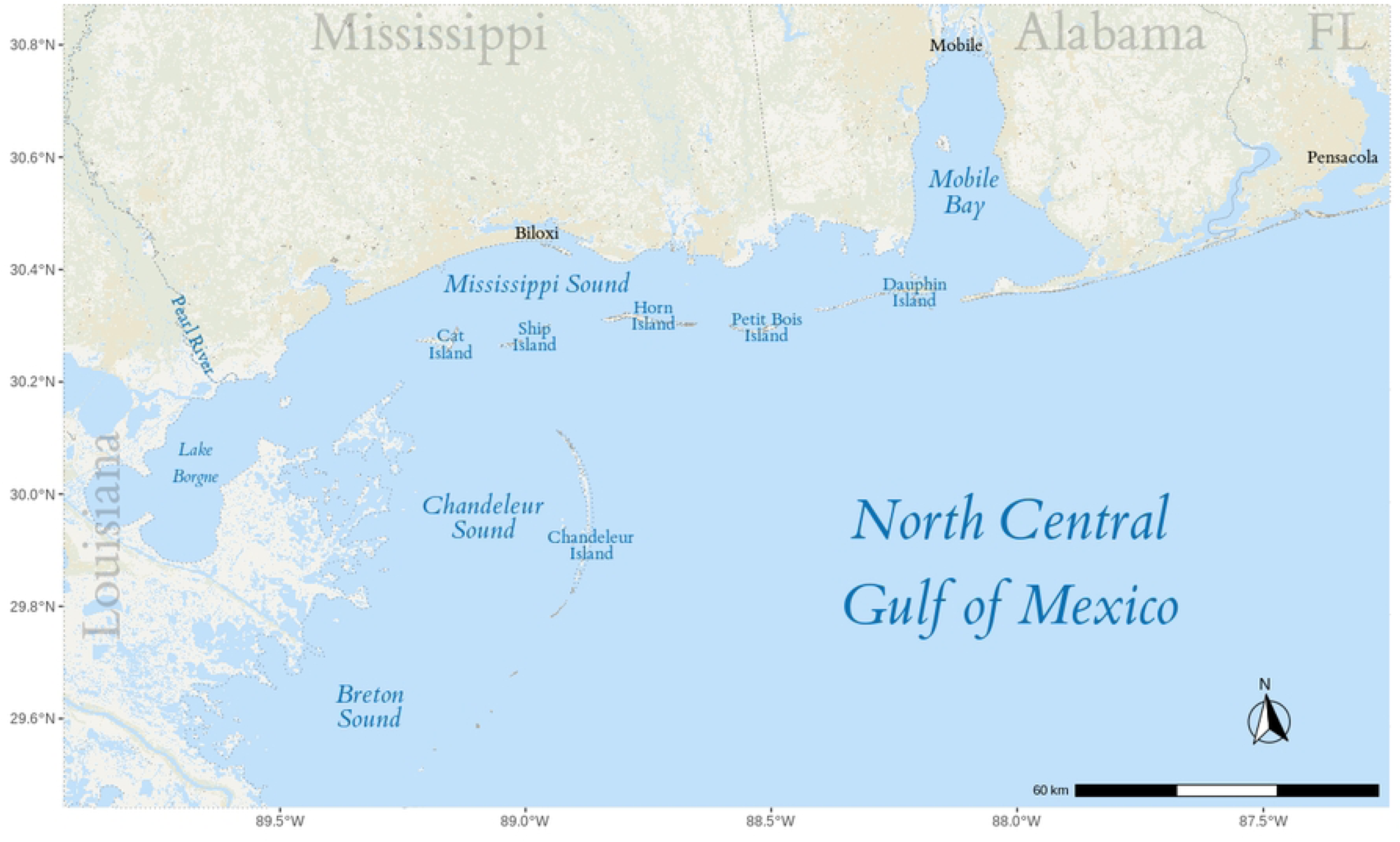
Map of sampled area. Map of the North Central Gulf of Mexico and the adjacent contiguous bodies of water.

Analysis of stranded marine mammal carcasses is an alternative to live-capture sampling which can be expensive to conduct, restricted to population sample size, frequency of sampling period, and the season the study is conducted. Utilization of carcasses can be a valuable tool to monitor populations in the wild throughout the year. However, there are several challenges in using stranded marine mammals. One challenge is the difficulty in determining the source population of stranded animals, as ocean currents and wind can cause them to drift, complicating efforts to infer characteristics of free-ranging populations; however, some efforts have been made to model these effects and estimate the origins[13–15]. Another challenge in collecting genetic samples from cetacean carcasses is that sampling is typically done opportunistically, without a predetermined sample size or research area. In addition, the complexity of population structures can further complicate efforts to identify stranded animal source populations[16]. Nevertheless, stranded cetaceans have been successfully used for molecular species identification[17,18], assessment of genetic diversity[19,20], and population genetic structure[21,22], as well as to estimate demographic parameters of species or populations[23].

On average, approximately 50 cases of dead dolphin strandings are reported annually in and around the Mississippi Sound[2], the majority being classified as moderately autolyzed carcasses (code 3; [24]). Compared to the inconsistent availability of live tissues or fresh dead carcasses, this consistent availability of material can facilitate surveys of genetic diversity and survivability within and among bottlenose dolphin populations across time and during environmental stressors (e.g., natural and man-made disasters, climate variability, populations disease prevalence impact). Because the mtDNA is highly abundant and resistant to degradation (relative to nuclear DNA), it offers a distinct advantage when using advanced autolytic tissues[25] by enabling the utilization of these valuable tissue resources.

The objective of the current study is to enhance our understanding of the genetic groupings of common bottlenose dolphins in the Mississippi Sound. Here we use whole mitochondrial genome sequences (mitogenomes) to investigate genetic variability among common bottlenose dolphins stranded on the Mississippi coast between 2010 and 2021. Despite the aforementioned caveats to using stranded animals, this strategy allows us to sample yearly and consistently without the large effort and expense required by live-capture sampling methods, which also result in stress on these protected animals. We use the increased resolution of whole mitogenomes to improve our understanding of population structure within the Mississippi Sound.

## Materials and Methods

### Sample Collection

Common bottlenose dolphin tissues were collected from stranded dead dolphins under the authority of the Marine Mammal Stranding Agreement between the National Marine Fisheries Service and the Institute for Marine Mammals Studies (IMMS). Therefore, ethics approval was not required. Dolphin tissues (327 muscle, 159 liver, 14 lung, 6 kidney, and 5 brain) were collected from 511 dolphins stranded in the Mississippi Sound from 2010 to 2021 by the IMMS. The samples used in this study were from archived frozen (-20°C) tissues collected during necropsies. After sampling, the tissues were preserved by dropping small tissue pieces into 95% ethanol, which were then transferred to the Institute for Genomics, Biocomputing, & Biotechnology (IGBB) at Mississippi State University for analysis.

### Primer Design

All complete mitogenomes for marine dolphins (family Delphinidae) available from NCBI RefSeq (Table S1) were aligned against the complete common bottlenose dolphin (*T. truncatus*) mitogenome (NC_012059.1) using bwa fastmap (v0.7.10)[26] to find the set of longest pairwise exact matches. The coverage of all these sets of matches was calculated using bedtools (v2.25.0)[27]. Regions of the mitogenome that had exact matches longer than 23 base pairs (bp) across the aligned genomes were considered candidate locations for primers. Aided by Primer BLAST[28], these locations were manually screened, modified, and paired to produce candidate primer pairs (Table S2).

### Extraction, Amplification & Sequencing

Genomic DNA (gDNA) was extracted from the sampled tissues using the Qiagen DNeasy Blood & Tissue Kit (Qiagen, Germantown, MD, USA) following the manufacturer’s instructions. The quantity and purity of gDNA were assessed with a Nanodrop One spectrophotometer (Thermo Fisher Scientific, Waltham, MA, USA).

The complete mitogenome was amplified from extracted dolphin genomic DNA (20-30 ng) using four specific primer pairs (Table S2) with overlapping amplicon regions and Phusion Hot Start Flex Master Mix (New England Biolabs, NEB, Ipswich, MA, USA). After AMpure XP beads (Beckman Coulter, Indianapolis IN, USA) cleanup, the amplicon pools (four amplicons per pool per sample) were used for Nanopore barcoded amplicon library preparation with Nanopore Ligation Sequencing Kit (SQK LSK109; Oxford Nanopore Technologies, Oxford, UK) and Native Barcoding Expansion 96 Kit (EXP-NBD196; Oxford Nanopore Technologies, Oxford, UK) and sequenced on a GridION sequencer (Oxford Nanopore Technologies, Oxford, UK) using a Flongle flow cell (FLO-FLG001; Oxford Nanopore Technologies, Oxford, UK).

After each sequencing run, the raw data were aligned to the common bottlenose dolphin reference mitogenome using minimap2 (v2.17-r941)[29,30]. The base depth for primary alignments was calculated using samtools (v1.9.4)[31], and the median base depth for each sample was graphed using R (v4.0.2)[32] with the tidyverse (v1.3.1)[33] package. Amplicons for samples with complete mitochondrial amplification but low coverage (median base depth less than or equal to 15) were resequenced.

Once the sequencing was complete, variant calling was performed with clair3 (v0.1-r10)[34] using the amplicon reference sequences to identify likely heteroplasmic samples. Samples with more than two high-quality (QUAL ≥ 20, i.e. 99% probability that a variant exists at the site; and GQ ≥ 20, i.e. 1% probability the call is incorrect) heterozygous SNPs were discarded. Medaka (v1.5.0) (https://github.com/nanoporetech/medaka) was used to make a consensus sequence for each amplicon in each sample using amplicon sequences parsed from the common bottlenose dolphin reference mitogenome. The amplicon sequences for each sample were assembled using CAP3[35]. The complete mitogenomes were trimmed of overlap using merge, part of the EMBOSS tool suite (v6.6.0)[36], and rotated to start at the same base as the reference common bottlenose dolphin mitogenome using circlator (v1.5.5)[37]. To validate the likely species of each sample, the consensus sequence for each sample was aligned using BLAST+ (v2.14.0)[38] to the mitogenomes used during primer design (Table S1). Any sample whose best alignment was a species other than *T. truncatus* was removed from further analysis.

Previously, genetic populations of bottlenose dolphins inhabiting the Mississippi Sound and coastal waters of the north-central Gulf of Mexico[11] and, more broadly, the coastal and offshore waters of the northern Gulf of Mexico[39] were studied using a portion of the mitochondrial control region (mtCR), along with other methods. To compare our samples to these two studies, the medaka polished amplicons that contained the mtCR region (P6) were reduced to the region between the primer pairs used to generate the published data (i.e., L15824 and H16265 or H16498;[40,41]) using cutadapt (v4.6)[42], discarding any sample without a match to L15824.

### mtCR and Whole Mitogenome Analyses

The newly sequenced and previously published mtCR sequences (Table S3) along with two *Stenella frontalis* mtCR sequences (DQ060054 and DQ060057) were aligned using mafft (v7.471)[43]. The mafft alignments were trimmed to remove gaps at either end of the alignments and any ambiguous base was resolved using the distribution of bases across all alignments using the R package ape (v5.7-1)[44]. The R package haplotypes (v1.1.3.1)[45] was used to condense the sequences into haplotypes. A Neighbor-joining tree was created using the R package ape (v5.7-1)[44] and visualized using tidyverse (v2.0.0)[33], ggtree (v3.6.2)[46], and aplot (v0.2.1)[47].

Both the principal component analysis (PCA) and population structure prediction were run using LEA (v3.0.0)[48–51] for K=1 to K=15 using the haplotypes after removing the outgroups. The population structure was run with ten repetitions per K. The percentage of variance explained for the principal components was calculated using the Tracy-Widom test[50,52] and used to generate a scree plot. The scree plot, using Cattell’s rule[53], and the lowest average cross-entropy were used to choose the best K (number of ancestral populations) for further analysis. The run (i.e., repetition) with the lowest cross-entropy from the selected K was graphed using tidyverse and used to group samples based on the highest admixture coefficient (q).

The nucleotide diversity (π), haplotype diversity, and population differentiation (Fst) were calculated for the trimmed dataset with the assigned group using the R packages pegas (v1.2)[54] and heirfstat (v0.5-11)[55]. An additional measure of population differentiation, ΦST, along with private alleles, was calculated using the R (v4.2.2) package poppr (v2.9.4)[56,57] with the adegenet (v2.1.10)[58,59] and ape (v5.7-1)[44] packages.

Similar methods were used to analyze the whole mitogenome sequences, albeit without the inclusion of published data and without trimming the mafft alignments. The complete mitogenome of *S. frontalis* (NC_060612.1) was used as an outgroup for phylogenetic analysis. Additionally, because the number of individuals assigned to each group was vastly different, the large groups were randomly sub-sampled into partitions roughly equal to the smallest group to calculate pairwise Fst and ΦST. For the partitioned groups, the average statistic across all partitions is reported.

## Results

### Data Generation and Quality Control

Universal primer pairs were designed to amplify the mitogenome of marine dolphins (family Delphinidae) from 33 conserved regions ranging from 24 bp to 105 bp each. Ten candidate primer pairs were designed (P1 - P10; Table S2; Fig S1) and tested, resulting in the selection of four primer pairs (P4, P6, P7, and P10) as the best combination to retrieve the entire mitogenome of 16.3 Kbp.

Tissue samples from stranded dolphins were collected over 11 years, from 2010 to 2021, resulting in 511 samples. Although most gDNAs were degraded, the extractions provided suitable templates for mtDNA amplification. Of the 511 samples collected, 417 had at least part of the mitogenome amplified (384 complete; 33 partial amplifications; Table S4) from fresh dead (decomposition code 2; 66 individuals) to advanced decomposition (code 4; 33 individuals) samples, with most coming from moderately decomposed animals (code 3; 318 individuals). Notably, redundancy between primers P6, P7, and P10 (Fig S1) resulted in fully amplified mitogenomes in 6 of the 33 samples with partial amplification. To ensure the accuracy and reliability of our data, we implemented quality control procedures that required resequencing of individual amplicons or entire samples based on read depth and amplicon presence (see methods). In total, we identified 44 samples that required resequencing, with nine requiring multiple rounds to ensure data accuracy. The average median read coverage for the complete samples were 236.1, 322.1, 546.3, and 320.5 reads per sample for P4, P6, P7, and P10 primer pairs, respectively (Fig S2).

To test for heteroplasmy, heterozygous SNPs were identified for each sample, and any sample with more than two heterozygous SNPs was removed from further analysis. Thirty-eight samples had at least one heterozygous variant; however, four (SER10-0256, SER11-0942, SER11-1021, and SER13-0420) were removed from further analysis due to excessive heterozygosity (4 – 37 heterozygous SNPs).

The 413 remaining samples passing the above criteria were assembled, trimmed, and rotated. All complete samples, along with six full-mtDNA partial samples, were assembled into whole, circular mitogenomes (16,387-16,405 bp). Smaller assemblies were generated for 27 samples (6,104 – 14,627 bp) where the entire mitogenome was not amplified. Of those, SER12-0678 was the only sample to recover only disjointed amplicons (i.e., P4 and P6).

In addition, the partial mtCR region used in previous studies[11,39] was extracted from each sample to use in combined analyses with the previously published data. Since P6 encompasses most of the partial mtCR region used in previous studies (ending approximately ten bases before the primer H16232), the ten samples lacking P6 (SER11-0040, SER11-1421, SER11-2252, SER11-2358, SER11-2425, SER12-0271, SER12-0735, SER13-0635, SER16-00070, and SER19-00351) were removed from further analysis. The P6 amplicon sequence in the remaining 403 samples was trimmed to the forward primer used in previous studies (L15824), producing an approximate 421 bp sequence for each sample.

### mtCR Haplotypes

To compare our samples with those previously published, we first reconstructed mtCR haplotype using all samples combined. The 511 mtCR sequences (106 previously published, 403 newly sequenced, and 2 outgroups) were aligned using mafft (Fig S3) and trimmed to 354 bases, representing 74 (72 bottlenose dolphin, 2 outgroups) haplotypes (sizes ranging from 1-198 members; Table S3 and S4), using the haplotypes R package. Of the 72 bottlenose dolphin haplotypes (Table S5), 52 contain only published sequences (mtCR.pub), 12 contain both published and new sequences (mtCR.mix), and 8 contain only new sequences (mtCR.new). Almost half (190) of the new samples are contained within a single haplotype (mtCR.mix-1) and most (363) are contained within the top five haplotypes. Most of the mtCR.new groups are composed of single individuals (six of eight), with the remaining two (mtCR.new-1 and mtCR.new-2) containing three and two samples, respectively. Among the mtCR.new groups, all differ from an mtCR.mix group by a single nucleotide (Fig S4).

A neighbor-joining tree was created using the 74 haplotypes (Fig 2A). After removing the outgroup sequences, LEA was used for the principal component and population structure analyses, evaluating up to 15 ancestral populations (K=1 to K=15) using the 44 biallelic SNPs present in the 72 haplotypes. Both the PCA scree plot (Fig S5A) and the cross-entropy plot (Fig S5B) suggest that four ancestral populations (K=4) best describe the data (Fig 2B). Each haplotype was assigned a group based on the highest admixture coefficient (q). Additionally, the published samples assigned to populations described in Vollmer and Rosel[39] and Vollmer et al.[11] were linked to the current haplotypes (Fig 2C). As shown in Table S6, the largest population (mtCR.sound) contains almost all (99.5%) newly sequence samples, as well as the majority (88%) of the previously described Northwest inner[39] and almost all (97%) Green (inshore and coastal waters west of Mobile Bay)[11] population sequences. The samples for the smaller Blue population (coastal waters east of Mobile Bay)[11] are evenly split between the mtCR.inner (47%) and mtCR.outer (44%) populations and is largely absent from both mtCR.sound (9%) and mtCR.ocean (0%). In the context of the previously described Vollmer and Rosel[39] populations, mtCR.inner contains a majority (55%) of the East inner and some (12%) Northwest inner sequences, and mtCR.outer contains a majority (61%) of the East outer and Northwest outer sequences, as well as more than half (57%) of Northwest oceanic sequences. The mtCR.ocean group largely consists of oceanic populations from Northeast, East, and Northwest. Taken together, the structure appears to transition from Green/NW inner (mtCR.sound), to Blue/inner (mtCR.inner), then Blue/outer/NW oceanic (mtCR.outer), and finally oceanic (mtCR.ocean) populations. The member haplotypes of each group are largely congruent with the populations described in both Vollmer et al.[11] and Vollmer and Rosel[39]; however, since the structure analysis did not include the microsatellite or SNP data some of the populations found in Vollmer et al.[11] and Vollmer and Rosel[39] were combined.

**Fig 2.**
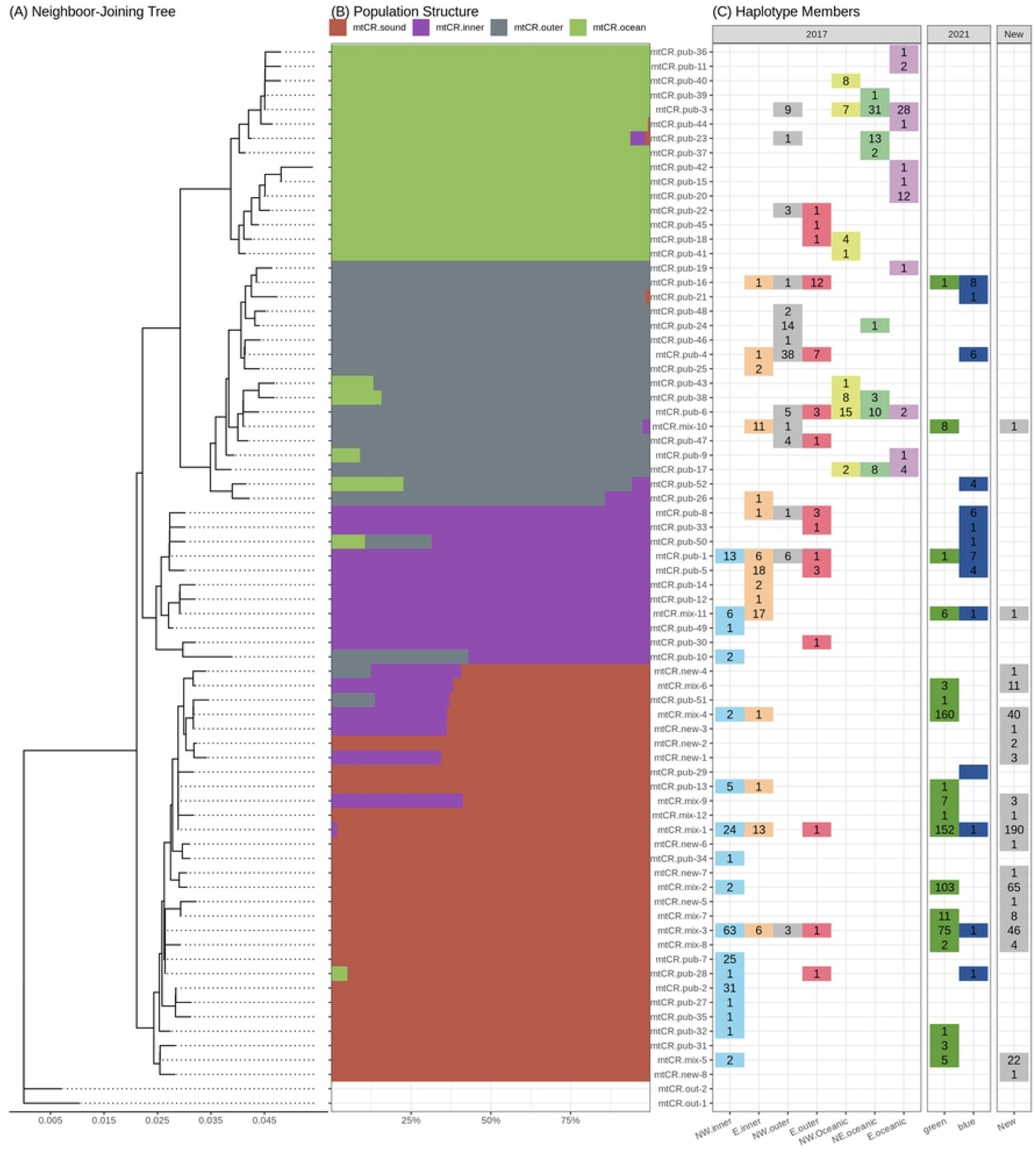
mtCR analyses. (A) Neighbor-joining tree, (B) structural analysis (K=4), and (C) the number of samples in each haplotype assigned to the populations described in Vollmer and Rosel[39] and Vollmer et al.[11].

Overall haplotype diversity was 0.8083 (±2.012x10^-4^) among these samples, with haplotype diversity for each group ranging from 0.7562 (±2.671x10^-4^) in mtCR.sound to 0.9779 (±7.426x10^-4^) in mtCR.ocean. The overall haplotype diversity falls within the ranges reported in Vollmer and Rosel[39] (0.6916 – 0.8633) and Vollmer et al.[11] (0.7816 – 0.8943); however, the per-group haplotype diversity range extends higher. Overall nucleotide diversity (π) was 0.0113 (±3.939x10^-5^), and the per group nucleotide diversity ranges from 0.0037 (±6.681x10^-6^) for mtCR.sound and 0.0087 (±2.818x10^-5^) for mtCR.ocean, on the lower end of the ranges reported in Vollmer and Rosel[39] (0.0062 – 0.0200) and Vollmer et al.[11] (0.0053 – 0.0202). The overall differentiation (Fst) was 0.0627 using the Weir and Cockerham method[60], with pairwise Fst (Table S7) ranging from 0.0296 (mtCR.inner vs mtCR.sound) to 0.0867 (mtCR.sound vs mtCR.ocean). These values are lower than the values reported in Vollmer and Rosel[39] (0.18 overall Fst) and Vollmer et al.[11] (0.16 overall Fst); however, this is not unexpected since the populations inferred using LEA combined several of the published populations. The overall ΦST was 0.6181, falling between the values reported in Vollmer and Rosel[39] (0.55) and Vollmer et al. [11] (0.65). The pairwise ΦST (Table S7) ranged from 0.4239 (mtCR.inner vs mtCR.outer) to 0.7442 (mtCR.ocean vs mtCR.sound), which is higher than reported in Vollmer et al.[11] (0.36 – 0.51), but within the range in Vollmer and Rosel[39] (0.041 – 0.799). All four groups have private alleles, ranging from five for mtCR.inner to nine in mtCR.sound.

### Full mitogenome

Because the full mitogenome may offer additional resolution[61–63], the 386 complete mitogenomes and the outgroup mitogenome from *S. frontalis* were aligned using mafft, resulting in an alignment length of 16,508 bases. These 386 common bottlenose dolphin sequences were condensed into 117 haplotypes, most (84) of which comprise single individuals. As expected, these haplotypes are consistent with the mtCR haplotypes but with additional resolution (Fig S6).

PCA and putative population structure were analyzed (via LEA) using 309 biallelic SNPs identified among the 117 haplotypes. The PCA scree plot (Fig S7A) and the minimum cross-entropy plot (Fig S7B) suggest that the data represent four populations (K=4, hereafter mitogroups 1-4; Fig 3A). As with the mtCR region, each haplotype was assigned to a mitogroup based on the maximum admixture coefficient (q), clustering 237 samples (67 haplotypes) in mitogroup 1; 35 samples (14 haplotypes) in mitogroup 2; 113 samples (35 haplotypes) in mitogroup 3; and 1 sample (1 haplotype) in mitogroup 4. The neighbor-joining tree (Fig S8) is largely congruent with the structure analysis. Given the admixture coefficients (Table S8)of the included samples show two distinct subgroups in mitogroup 3 , it is not unsurprising that this group is paraphyletic; which could suggest a large ancestral population that has separated over time.

**Fig 3.**
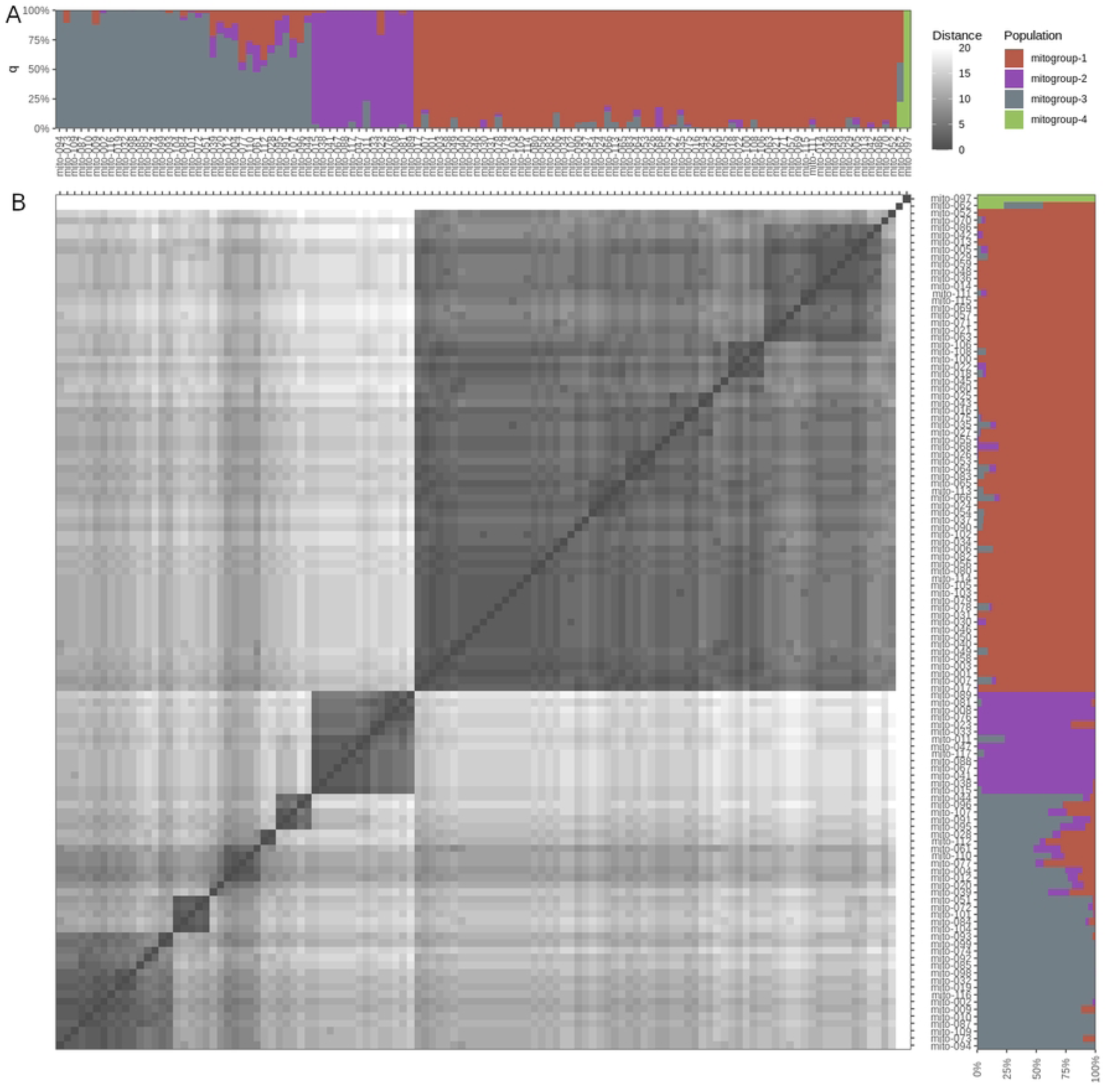
Mitogenome population structure. (A) LEA structure analysis (K=4) and (B) sequence similarity of the whole mitogenomes.

Only two haplotypes (mito-097, q = 0.9997, and mito-062, q = 0.2250) had an admixture coefficient in mtiogroup 4, with mito-097 being the only haplotype assigned to that mitogroup. Notably, the two sequences represented by these two haplotypes are the only two samples not assigned to the mtCR.sound group in the mtCR analysis; i.e., SER11-0141 was assigned to mtCR.inner and SER19-00888 to mtCR.outer. Additionally, these two haplotypes are unusually dissimilar compared to the other sequences (Fig 3B) and, in the neighbor-joining tree, these two haplotypes are separated from the other clades (Fig S8). Taken together, these two samples likely originated from population outside the sampled area. Since mitogroup 4 only contained a single haplotype, it was dropped from most downstream analyses.

The stranding locations per group are shown in (Fig 4). The stranding location areas of group 2 and 3 are contained within that of group 1, with group 1 extending as far east as Pensacola, FL and as far west as Lake Borgne. Interestingly, the stranding region of group 2 is noticeably smaller than that of both group 1 and 3, extending from the Pearl River to Biloxi, MS. To test if the smaller region is due to the smaller population size, a bootstrap test with 1,000,000 iterations was run against the mean longitude using the boot (v1.3-30)[64,65] and boot.pval (v0.5)[66] R libraries. P-values were calculated using the normal approximation confidence intervals. The mean longitude of groups 1 and 2 are significantly different (p-values: 0.00437 and < 10^-6^, respectively) from the entire dataset.

**Fig 4.**
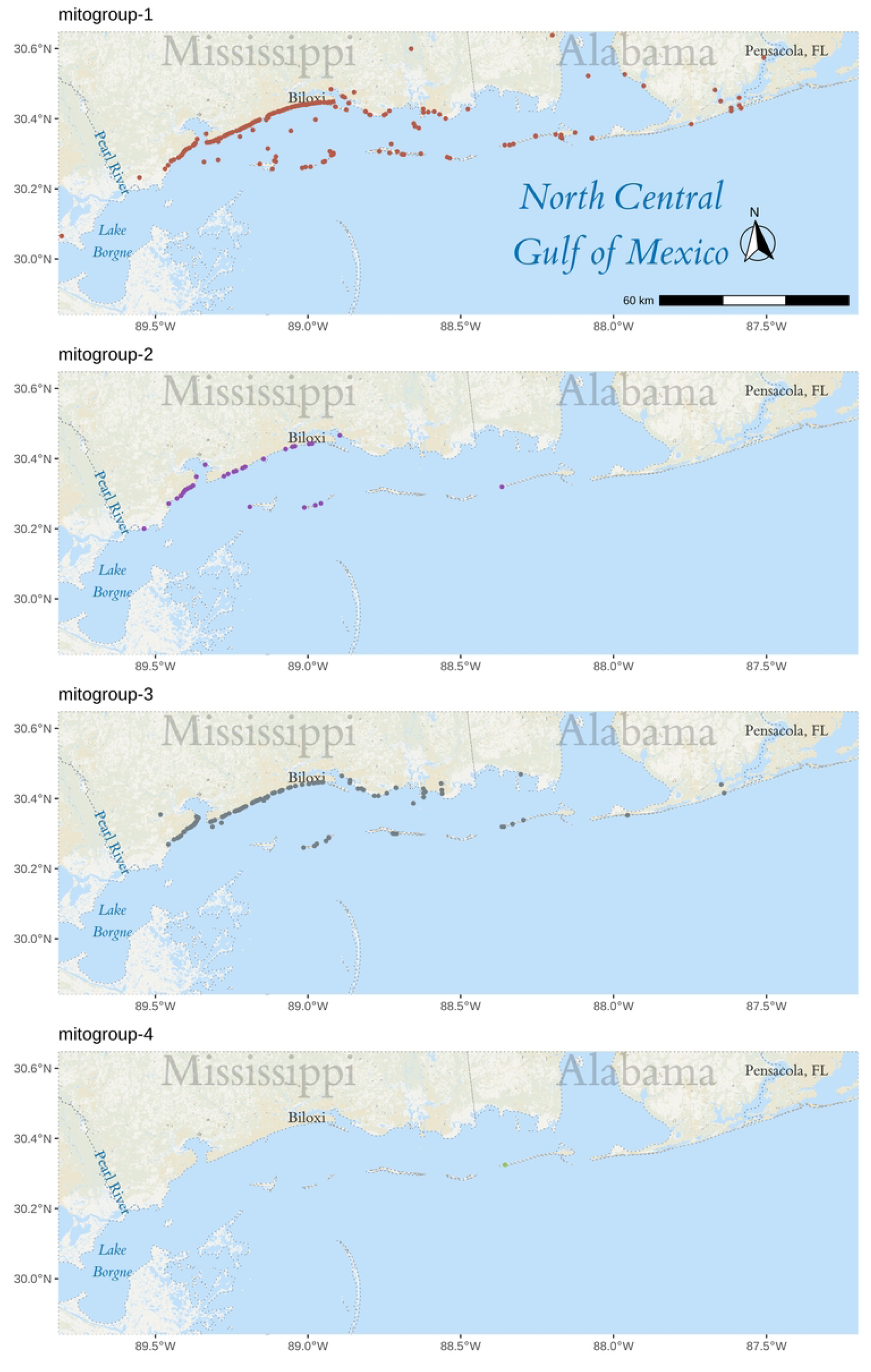
Stranding locations per mitogroup. Map of the stranding location for each group. Map created used the tidyverse (v1.2.1)[33], sf (v1.0-18)[82,83], ggspatial (v1.1.9)[84], and stars (v0.6-6)[83] R libraries with data from the USGS 100-Meter Resolution Land Cover of the Conterminous United States.

The overall haplotype diversity was 0.9363 (±6.34x10^-5^), while the individual group haplotype diversity was 0.8604 (±3.70x10^-4^), 0.8857 (±8.46x10^-4^), and 0.8761 (±4.24x10^-4^) for mitogroups 1 through 3, respectively. The nucleotide diversity (π) was 0.00051 (±6.77x10^-8^) overall and 0.00017 (±1.01x10^-8^), 0.00018 (±1.15x10^-8^), and 0.00031 (±2.80x10^-8^) for mitogroups 1, 2, and 3, respectively. Since both mitogroups 1 and 3 are much larger than group 2, the pairwise population differentiation statistics Fst and ΦST were calculated on 6 and 3 subsets, respectively, and averaged for each comparison that included either of those two groups. Overall Fst and ΦST were 0.5784 and 0.7250, respectively. Pairwise differentiation ranges from 0.6475 (between mitogroups 1 and 3) to 0.8612 (mitogroups 2 and 4) for Fst and 0.6646 (between 1 and 3) and 0.8612 (between 2 and 4) for ΦST. Only mitogroup 1 had any private alleles.

## Discussion

Genetic diversity is an important factor in the long-term survival of any given population. While cetaceans are highly mobile animals with a high potential for dispersal, several studies have found a range of impediments to gene flow resulting in population sub-structuring in some species[67–69]. Therefore, understanding the genetic diversity and the population structure of common bottlenose dolphins is vital to conservation, monitoring, and marine management projects. To this end, there is an ever-increasing interest in applying molecular techniques such as single nucleotide polymorphisms (SNPs), mitochondrial DNA (mtDNA), nuclear microsatellites, and more recently, environmental DNA (eDNA), for identifying and monitoring populations of dolphins[70–75] for conservation purposes.

The aim of this study was to investigate the genetic variability of common bottlenose dolphins stranded on the Mississippi coast from 2010-2021. The tissue samples used in our study were collected during necropsies of stranded dolphins, which presented a challenge to isolate high-quality DNA due to the varying degrees of decomposition. Thus, we decided to amplify and use the whole mitogenome sequence. Most DNA isolations were from muscle and liver tissues, which are expected to have high and medium mitochondria numbers due to their respiratory activities[76]. We used four primer sets to successfully amplify 417 mitogenomes out of 511 DNA samples (81.29% success rate), 386 (92.57%) of which resulted in full mtDNA amplification. Using the assembly methods described, the overlapping region added by P10 is superfluous and can be omitted. We obtained complete mitogenomes from stranded samples ranging from fresh dead (code 2) to advanced decomposition (code 4), with most of the samples being moderately decomposed (code 3). Interestingly, the decomposition does not seem to affect the likelihood of successful mtDNA amplification. We excluded mummified (code 5) remains; however, with current method advancements[77] in hard tissue DNA extraction, those samples could be an additional source of mtDNA to consider.

It is worth noting that our study used a different methodology than Vollmer et al.[11] and Vollmer and Rosel[39]. These two previous studies used skin samples from live common bottlenose dolphins and a combination of nuclear genomic sources and part of the mitochondrial control-region (mtCR). Since the newly sequenced samples were collected from stranded dolphins at several stages of decomposition, the extracted DNA was often highly degraded and contained high levels of bacterial contamination. While it is possible to analyze small allele microsatellites (<200 bp) using highly degraded DNA[78], some of the microsatellites targeted in previous studies do not fit into this category. Since it was likely we would not be able to use some of the microsatellites used in previous studies due to the decomposition level of many samples, we focused on amplifying the entire mitogenome. By using the entire mitogenome, we increased the number of usable SNPs identified (309 vs 44), allowing for more precision in identifying populations (the single population containing almost all newly sequenced haplotypes in Fig 2B vs the three groups in Fig 3A), while simultaneously reducing the impact of homoplasy, which is common in the mtCR.

Previous research in marine animals has demonstrated the value in using mitogenomes to analyze complex evolutionary relationships. For example, Leslie, Archer, and Morin[61] used whole mitogenome sequences from spinner (*Stenella longirostris*) and pantropical spotted dolphins (*Stenella atteunuata*) to test population structure hypotheses at multiple hierarchical taxonomic levels, although the ability to distinguish among populations units within each species was interestingly not improved from earlier studies by the inclusion of the entire mitogenome. Conversely, other investigators[62] used next-generation sequencing (NGS) to generate a total of over 500 whole mitogenomes from seven cetacean species and green sea turtles, showing that entire mitogenomes exhibited greater variation than shorter regions, thereby improving resolution. Likewise, Nykänen et al.[63] found entire mitogenomes improved resolution in reconstructing post-glacial expansion of the Northeast Atlantic bottlenose dolphins. While these and other studies clearly demonstrate the utility of mitogenomes in providing insight into marine populations, it is important to note that, although mtDNA sequencing can provide valuable insight into genetic diversity and migration, it only provides information about the maternal lineage and may not reflect the overall genetic diversity of a population, and our study would benefit from additional nuclear genomic sources. Given the decreasing cost and improving quality of long-read sequencing technologies, future studies will benefit from scaling up to whole mitogenome as well as including nuclear markers, such as SNPs or microsatellites. Additionally, given the large overlap between the four amplicons, it may be possible to generate and include the multiple genomes of heteroplasmic samples in the structural and comparative analyses.

Population structure analysis and differentiation statistics were run twice: once with the mtCR region of the stranded samples and published data used in Vollmer and Rosel[39] and Vollmer et al.[11], and again using the whole mitogenome for the stranded samples that could be completely assembled. The mtCR analysis suggests a single group that commonly strands along the Mississippi coast, corresponding to the green population in Vollmer et al.[11] and the Northwest inner population in Vollmer and Rosel[39], with two samples likely originating outside the Mississippi Sound. By comparison, whole mitogenome structure analysis and differentiation statistics also suggest two samples originated outside the sample area, but this method defines a slightly more complex system for the Mississippi Sound, with two groups that commonly strand across the entire sampled region (mitogroup 1 and 3) and a group (mitogroup 2) that occasionally strands in a specific subset (between the Biloxi Bay and Pearl River). Some species of marine mammals show genetic differentiation over small geographic scales, despite the fact that they are generally considered to have a high propensity to disperse; e.g., Indo-Pacific bottlenose dolphins (*T. aduncus*)[79], Commerson’s dolphins (*Cephalorhynchus commersonii*)[80], and harbor porpoises (*Phocoena phocoena*)[81].

While genetic research based on carcass samples may be difficult to interpret due to the disjunction between the actual habitat utilized while alive and the post-mortem stranding location, the latter of which is influenced by factors such as sea currents and the unpredictable behavior of dying dolphins, our results suggest that this can be a valuable sampling strategy for restricted or difficult to access populations, such as the common bottlenose dolphin. Our findings align with the population structure reported previously[11,39]. The quantity of mitochondrial diversity recovered in the present study suggests there are different subpopulations of common bottlenose dolphins in the Mississippi Sound area, providing finer-scale resolution of genetic structure among these mammals than previously reported.

Our research demonstrates that whole mitogenome sequences derived from various tissues sampled from stranded carcasses, with muscle and liver being best, may be suitable for characterizing and monitoring the genetic diversity of cetaceans in areas like the Mississippi Sound. It also provides important insights into the genetic diversity of common bottlenose dolphins in the Mississippi Sound, which can help inform conservation efforts and management decisions for this population.

## Acknowledgments

We thank IMMS for providing frozen dolphin tissues collected during necropsies of stranded dolphins. Funding for this project was generously provided by the Mississippi Department of Marine Resources (Gulf of Mexico Energy Security Act project number 3000027046).

## DATA AVAILABILITY

All data are available at NCBI under the BioProject PRJNA842356 and under the SRA (Sequence Read Archive) accessions SRR19396654 - SRR19397072. The mitochondrial genomes are available from GenBank (OR612358 - OR612776). The code used for analysis is available at https://github.com/IGBB/ms-sound-dolphin.

## Captions

### Supplementary Tables

S1 Accession number and species of mitochondrial genomes used in primer design https://docs.google.com/spreadsheets/d/1XtFSuL4BN1HTnHtHo3o0YAuOiTaBu8MWYjtfvLf 0ltc/edit?gid=1574452535#gid=1574452535

S2 Tested primer pairs, highlighted pairs were used in final analysis https://docs.google.com/spreadsheets/d/1XtFSuL4BN1HTnHtHo3o0YAuOiTaBu8MWYjtfvLf0ltc/edit?gid=0#gid=0

S3 Previously sequenced mtCR haplotypes of GoMx bottlenose dolphins with the new haplotype and assigned group, along with the population assignments in Vollmer and Rosel[39] and Vollmer et al[11] https://docs.google.com/spreadsheets/d/1XtFSuL4BN1HTnHtHo3o0YAuOiTaBu8MWYjtfvLf0ltc/edit?gid=1299223648#gid=1299223648

S4 Information of sequenced samples from this study https://docs.google.com/spreadsheets/d/1XtFSuL4BN1HTnHtHo3o0YAuOiTaBu8MWYjtfvLf0ltc/edit?gid=123140332#gid=123140332

S5 Summary of mtCR haplotypes https://docs.google.com/spreadsheets/d/1XtFSuL4BN1HTnHtHo3o0YAuOiTaBu8MWYjtfvLf0ltc/edit?gid=1500357613#gid=1500357613

S6 Comparison of membership sample and haplotype totals of each mtCR inferred population and the reported population in Vollmer and Rosel[39] and Vollmer et al.[11] https://docs.google.com/spreadsheets/d/1XtFSuL4BN1HTnHtHo3o0YAuOiTaBu8MWYjtfvLf0ltc/edit?gid=2142620833#gid=2142620833

S7 Population Statistics https://docs.google.com/spreadsheets/d/1XtFSuL4BN1HTnHtHo3o0YAuOiTaBu8MWYjtfvLf0ltc/edit?gid=2112999263#gid=2112999263

S8 Full mitogenome LEA admixture for the run with the lowest cross entropy using K=4 https://docs.google.com/spreadsheets/d/1XtFSuL4BN1HTnHtHo3o0YAuOiTaBu8MWYjtfvLf0ltc/edit?gid=1629546585#gid=1629546585

### Supplementary Figures

S1 Tested primer pair locations on the mitochondrial genome. Highlighted pairs (P4, P6, P7, and P10) were used to amplify the mtDNA in all samples. https://github.com/IGBB/ms-sound-dolphin/blob/master/1-primer-design/primer-locations.png

S2 Median coverage for each amplicons https://github.com/IGBB/ms-sound-dolphin/blob/master/2-samples/amp-coverage.png

S3 Alignment of all mtCR sequences (both published and newly sequenced). The missing data (black) at both ends were trimmed. https://github.com/IGBB/ms-sound-dolphin/blob/master/4-mtCR/alignment.outliers.png

S4 Nucleotide differences between the mtCR.new haplotypes and the most similar sequences. All sequences with one or fewer mismatched bases to the haplotype of interest are displayed. All identical bases are removed for clarity. https://github.com/IGBB/ms-sound-dolphin/blob/master/4-mtCR/unique-haplotypes.png

S5 PCA Scree plot (A) and Minimum cross-entropy per K (B) for the mtCR structure analysis https://github.com/IGBB/ms-sound-dolphin/blob/master/4-mtCR/4-struct-fig-K-fixed.png

S6 Sankey plot showing the change in haplotypes from mtCR to whole mitogenome. Haplotypes with five or more members are labeled. https://github.com/IGBB/ms-sound-dolphin/blob/master/5-full-mito/mtCR-vs-whole-sankey.png

S7 PCA Scree plot (A) and Minimum cross-entropy per K (B) for the whole mitogenome structure analysis https://github.com/IGBB/ms-sound-dolphin/blob/master/5-full-mito/3-structure-haplotype-fig-K.png

S8 Neighbor-joining tree for the whole mitogenome. Outgroup is the *S. frontalis* sequence NC_060612.1. Color shoe the assigned group of each sample. https://github.com/IGBB/ms-sound-dolphin/blob/master/5-full-mito/full-mito-nj-tree-circular.png

